# Accurate Automatic Glioma Segmentation in Brain MRI images Based on CapsNet

**DOI:** 10.1101/2021.07.03.450931

**Authors:** M. Jalili Aziz, A. Amiri Tehrani Zade, P. Farnia, M. Alimohamadi, B. Makkiabadi, A. Ahmadian, J. Alirezaie

## Abstract

Glioma is a highly invasive type of brain tumor with an irregular morphology and blurred infiltrative borders that may affect different parts of the brain. Therefore, it is a challenging task to identify the exact boundaries of the tumor in an MR image. In recent years, deep learning-based Convolutional Neural Networks (CNNs) have gained popularity in the field of image processing and have been utilized for accurate image segmentation in medical applications. However, due to the inherent constraints of CNNs, tens of thousands of images are required for training, and collecting and annotating such a large number of images poses a serious challenge for their practical implementation. Here, for the first time, we have optimized a network based on the capsule neural network called SegCaps, to achieve accurate glioma segmentation on MR images. We have compared our results with a similar experiment conducted using the commonly utilized U-Net. Both experiments were performed on the BraTS2020 challenging dataset. For U-Net, network training was performed on the entire dataset, whereas a subset containing only 20% of the whole dataset was used for the SegCaps. To evaluate the results of our proposed method, the Dice Similarity Coefficient (DSC) was used. SegCaps and U-Net reached DSC of 87.96% and 85.56% on glioma tumor core segmentation, respectively. The SegCaps uses convolutional layers as the basic components and has the intrinsic capability to generalize novel viewpoints. The network learns the spatial relationship between features using dynamic routing of capsules. These capabilities of the capsule neural network have led to a 3% improvement in results of glioma segmentation with fewer data while it contains 95.4% fewer parameters than U-Net.

## Introduction

Gliomas are the most common fatal brain tumors caused by the abnormal growth of glial cells in the brain [1, 2]. It is including different sub-regions, i.e., pre-tumoral edema, necrotic core, enhancing and non-enhancing tumor core. Among the various medical imaging techniques, Magnetic Resonance Imaging (MRI) has become the gold standard for brain tumor diagnosis due to its ability to provide good contrast for soft tissue and the availability of a variety of additional modalities [3, 4]. A clear appearance of the glioma sub-regions, can be found in four MRI sequences, including: T1-weighted (T1), T2-weighted (T2), T1-weighted with gadolinium contrast enhancement (T1-Gd) and Fluid attenuated Inversion Recovery (FLAIR) [5]. Gliomas can appear in any part of the brain and are heterogeneous in shape, size, and appearance with blurred and irregular borders that make identifying the exact boundaries in the image exceedingly difficult [6–8].

The maximal resection of the tumor while preserving the neighboring functional non-tumoral tissue is a pivotal goal in neuro-oncological surgery. Therefore image-guided surgery is nowadays an important part of the modern neuro-oncology practice. Tumor segmentation is an essential step in surgery under image-guidance, where the image visualization and registration depend on the accurate segmentation of the tumoral tissue [9–11]. Over time, different conventional brain image segmentation methods have been developed, including manual segmentation, intensity-based methods [12], surface-based methods [13], and other deformable models. In the manual segmentation method, the segmentation process is performed by trained clinicians based on their skills. Therefore, its results depend on the user’s subjective decision. Also, the time-consuming nature of manual segmentation is another limitation of these methods. However, manual segmentation is still necessary as a gold standard approach to evaluate other methods [5]. Despite many efforts that have been made to overcome the limitations of tumor segmentation algorithms, due to the highly heterogeneous nature of gliomas, conventional solutions have not been satisfactory. On the other hand, deep learning approaches has received a great deal of attention for many image processing applications such as image de-noising [14], image segmentation [15, 16], and image reconstruction [17].

Convolutional Neural Networks (CNNs), as one of the most successful deep learning methods, can provide accurate image segmentation results [18]. In this method, the network is able to learn useful features automatically, without the need for manual feature selection. CNNs have been widely used in the literature, with different approaches and have provided acceptable results in the brain tumor segmentation [5, 19–21]. Despite the recent significant achievements reported in the literature, there are four main shortcomings associated with CNNs. The first limitation is that CNNs cannot maintain the dependencies between the object parts and their totality, because of their structure design [22]. Second, the pooling layers use a type of routing which is not based on the human visual system. It routs important information which is extracted from the image to all the neurons of the next layer completely, thus details or small objects in the image are missed. The third limitation is that, in the pooling layer, the information is routed statically from one layer to the next, and the next layer of neurons is selected with no intuition. In comparison, in the human visual system, this is done dynamically and neurons from the next layer can choose what information is important [23]. The fourth and most serious issue with CNNs is the amount of data necessary for training. Tens of thousands of images are required for CNN training, and preparing huge datasets for medical applications is a challenging task [24].

To overcome these limitations of CNNs, a novel class of neural networks was proposed by Sabour et al [25], where a group of neurons represents the existence of features. This group of neurons is called a “Capsule”, and the network made up of these blocks is called a “Capsule Neural Network” (also called CapsNets). The vertices on a CNN network are neurons, which have a scalar representation of the output. In CapsNet, an advanced technique is used to connect the capsules between layers, which obtains the network weights based on an iterative optimization strategy. In this routing technique, the output of the previous capsule is given as input to the next capsule and during an iterative process, the similarity between the input and output of the capsule is compared. Finally, the previous capsules are routed to the capsule which has a similar output [25]. Recently, CapsNets have been used for medical image segmentation [26].

To the best of our knowledge, this is the first time that CapsNet has been optimized for glioma segmentation on MR images. In our proposed method, the network is able to train with a smaller dataset in comparison with the commonly used CNNs. This network, by using “routing by agreement”, detects the relationship between parts of an object and the whole object by performing an iterative process.

## method and material

### A. Dataset Description

Brain Tumor Segmentation (BraTS) is a well-accepted benchmark for the automatic brain tumor segmentation in multimodal MRI scans of high-grade and low-grade glioma. [27]. BraTS2020 which is used in this experiment, contains MRI scans of 369 patients with ground truth in four modalities (T1, T2, T1-Gd and FLAIR).

### B. Capsule Network and routing by agreement algorithm

CapsNet is a network that identifies the spatial and hierarchical relationships between objects in images. The network is composed of several capsule layers, which are trained with an iterative algorithm. Each capsule is made up of a number of neurons and its output is a vector, which is a richer representation of the output. This vector has two general characteristics, including length and orientation. The probability that the entity indicated by the capsule corresponds to the current input is represented by the vector length. The orientation of the vector indicates the state of an object. A simple capsule structure is shown in Fig. 1. Here these weights are determined through routing by agreement iterative algorithm [25]. A temporary variable (*b_ij_*) is established for updating the weights in this procedure, and it is set to zero at the start of the loop.

**Fig. 1.**
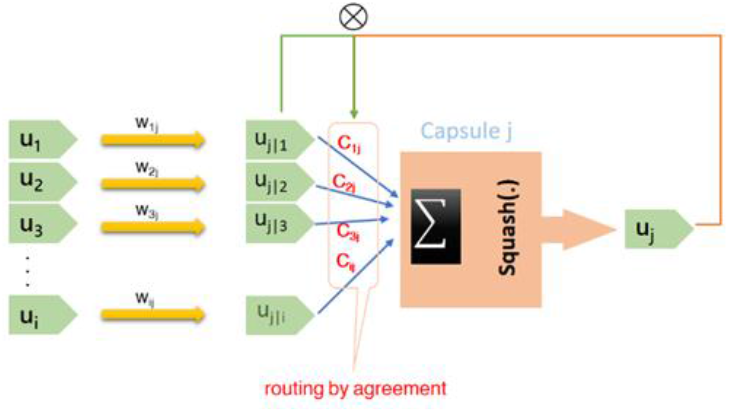
Capsule structure including inputs (*û_i_*) and output (*û_j_*).

In each iteration (*r*) for each capsule (*i*) in the layer *l*, this variable will be updated. In Fig. 1, *u*_0_,*u*_1_,*u*_2_, …, *u_i_* represent the output vectors of previous layer capsules (layer *l*) that encode the existence and the state of features in low level objects. Prediction vectors (*û*_*j*|*i*_) were obtained by multiplying *û_i_* by the corresponding weights. These weights show the relationship between the features obtained from lower-level capsules and higher-level capsules. *C_ij_* is a scalar weighing of inputs, called “coupling coefficient”, stored in *C_ij_* after passing the soft-max function. Initially, all routing weights are equal, this denotes the greatest degree of uncertainty in routing the previous layer capsules’ output vectors. In the next step, for the capsule *j* in layer *l* + 1, the input vector is calculated as (1):

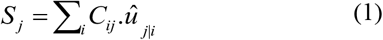

*S_j_* denotes the sum of weighted inputs. Next, for all capsules in layer *l* + 1, to determine capsule output vector (*V_j_*), *S_j_* is passed through a vector-to-vector function as below (2):

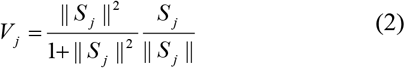

This nonlinear function compresses the vector’s length to the interval between zero and one, which is true in the sense that the output vector is probabilistic without affecting its direction. Then, for all capsules in the layer *l* and for all capsules in the layer *l* + 1, the temporary variable is updated as follows (3):

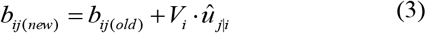

In this equation, the correspondence between the current output and *û*_*j*|*i*_, is shown by a dot product operation (Fig. 1). This routing algorithm calculates the output of the capsule *j* during an iterative process.

In this section, the basic concepts of capsule network architecture and its routing algorithm were stated. In the next section, we will describe the segmentation capsule network (SegCaps) architecture, which is an extension of the capsule network for segmentation tasks.

### C. SegCaps architecture

Due to the complexity of the CapsNet, the network encounters runtime and memory constraints for image segmentation, for the first time, LaLonde et al [28] introduced a new architecture called SegCaps based on CapsNet to address segmentation problems. In SegCaps framework, routing of lower layer capsules to the next layers is done only in a specific spatial window and the transformation matrixes are also shared between the capsules of each layer. The routing algorithm in SegCaps differs from CapsNet proposed in [25] in details.

*V_j_* in (2) is rewritten as *V*_*x*,*y*_; the output of the capsule at spatial coordinates (*x*, *y*) as follows (4):

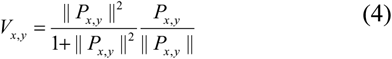

*P*_*x*,*y*_ is the capsule input. The agreement is defined as dot product of output vector and corresponding prediction vectors (5):

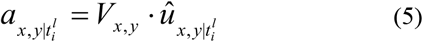

Also, in this architecture, a deeper network was introduced than CapsNet, by developing the concept of de-convolutional capsules. Therefore, the dimensions of the input image will be increased to 512 × 512, in which case the segmentation task is possible.

## Experiments and results

### A. Experimental Configuration

We implemented the proposed networks with Keras and TensorFlow framework using NVIDIA GeForce RTX 1080 TI GPU. All training is performed using dice loss and Adam optimizer. The data included MRI scans of 369 patients, of which 70 slices were used for each patient scan. Then 15% of this data is extracted randomly as a test dataset and the remaining 85% are used as training and validation set. We used the same test subset of the dataset to validate our results on SegCaps and U-Net. The image slices are cropped to 224× 224 and fed 4 channel images (including flair, T1, T1-gd, and T2) to network with random augmentation and the batch size of one. Implementation details and specific settings for each network are as follows:

- **SegCaps**: In this experiment, we randomly selected 20% of the total slices and trained the network in two steps. 80% of this subset is used for training and the remaining 20% is used as validation data. In the first phase, initially, we trained the network with a learning rate of 0.001, step decay of 1e-6 and reconstruction weight of 20. The weights of this phase are used as an initial weight for the second phase of training, via a learning rate of 1e-7 and reconstruction weight of zero.
- **U-Net**: We trained the U-Net network on 85% of the whole dataset as a training subset of the dataset. 80%of this subset is used as training and 20% as validation. The network is trained using a learning rate of 1e-7 and step decay of 1e-6 for 200 epochs.

### B. Analysis of Experiments

To evaluate our proposed approach quantitatively, BraTS2020 was used. Quantitative results are presented by the performance measurement in the form of Dice Similarity Coefficient (DSC) for validation data by U-Net and SegCaps in the Table.1. The DSC is calculated as (6) where P_1_ is the segmented region of the tumor and T_1_ is related area in the ground truth mask [3]:

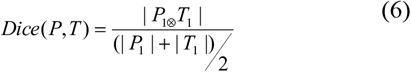

**TABLE I.** COMPARISION BETWEEN SEGCAPS AND U-NET ON BRATS2020 dataset

| Method | Parameters | DSC % |
| --- | --- | --- |
| U-Net | 31 M | 85.56 |
| SegCaps | 1.5M | 87.96 |

For qualitative evaluations, we have shown segmentation results using two approaches, U-Net and SegCaps, in comparison with Ground Truth, from three different patients with glioma tumors on multimodality MR images in Fig. 2. The three columns contain three patients’ MRI data, including T1, T1-Gd, FLAIR, and T2 sequences, in which the ground truth masks, U-Net, and SegCaps segmentation results are marked on the T2 images, respectively.

**Fig. 2.**
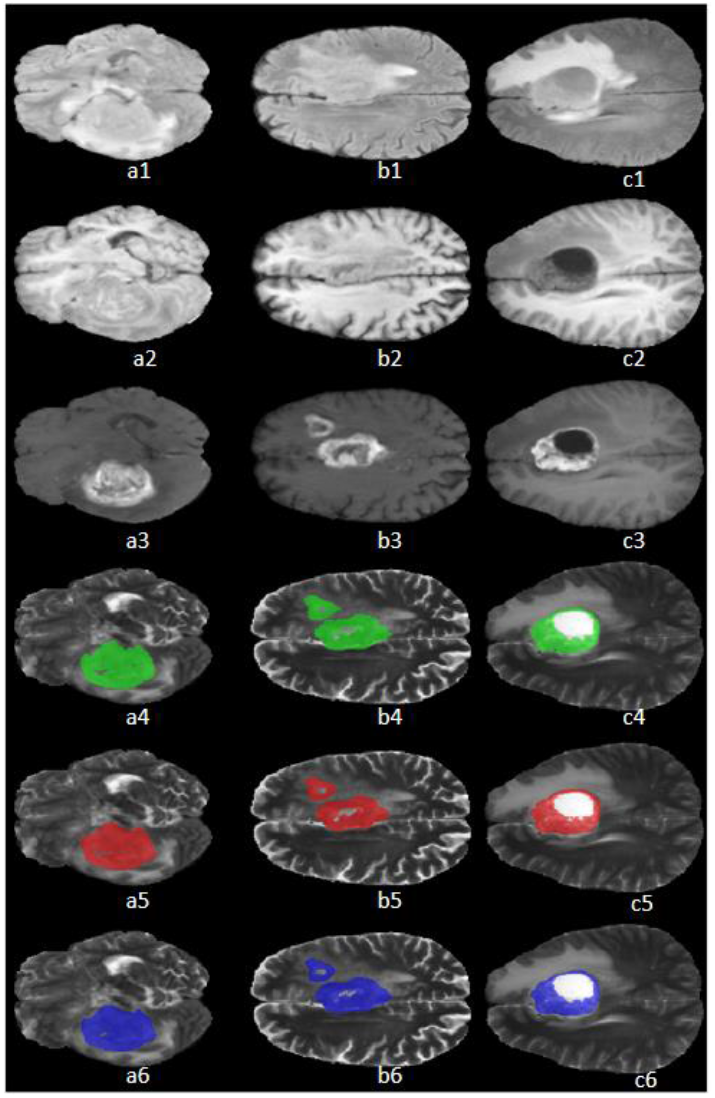
Qualitative results on the BraTs2020 dataset for three different patients: **a**, **b** and **c**. For each patient, from top to bottom: FLAIR, T1, T1-Gd, T2 with true mask (Green), T2 with U-Net mask (Red) and T2 with SegCaps mask (Blue), respectively.

## Conclusion

To achieve accurate glioma segmentation in MR images without the need for a huge number of training datasets, we have optimized the SegCaps architecture introduced by LaLonde et al. [36]. The results of our proposed approach are compared to the results of U-Net, a commonly used network for medical image segmentation. At first, we tried to train the SegCaps using the entire dataset. However, by using different hyper-parameters settings, the DSC did not improve more than 0.7 (70%) and the network could not converge due to the complexity of the BraTS dataset. Also, the complex architecture of capsules and routing algorithms make it very challenging to achieve convergence. Then, we used a subset of the dataset and trained the network in two steps. Since the capsule network has the intrinsic capability to generalize novel viewpoints, SegCaps learn the spatial relationship between features using dynamic routing of capsules. Therefore, it is expected that the SegCaps network, which is trained using a randomly selected subset of the dataset will have comparable results to the U-Net. As we have shown in Table. 1, by using the proposed two-step training method, our experimental results show that SegCaps has about 3% improvement compared to U-Net in DSC on validation data, while using fewer data for training and containing 95.4% fewer parameters than U-Net.

It can be concluded that the main advantage of the SegCaps is overcoming the problem of data amount limitation, which is a common issue in medical datasets. This can be qualitatively observed in Fig.2 that the SegCaps has been successful in segmenting the enhancing tumor core area. Despite capabilities of SegCaps, its routing algorithm is much slower than backpropagation. Also, its computational complexity, in addition to being time-consuming, will involve many GPU storage resources. However, since the SegCaps uses convolutional layers as a basic component, it has the potential to be optimized for complex segmentation tasks in challenging medical images, to achieve excellent results without data size limitation.

